# Origin, age, and metabolisms of dominant anammox bacteria in the global oxygen deficient zones

**DOI:** 10.1101/2023.10.31.564962

**Authors:** Rui Zhao, Irene H. Zhang, Amal Jayakumar, Bess B. Ward, Andrew R. Babbin

**Author notes:** Correspondence to RZ or ARB.

## Abstract

Anammox bacteria inhabiting oxygen deficient zones (ODZs) are a major functional group mediating fixed nitrogen loss and thus exerting a critical control on the nitrogen budget in the global ocean. However, the diversity, origin, and broad metabolisms of ODZ anammox bacteria remain unknown. Here we report two novel metagenome-assembled genomes of *Scalindua*, which represent most, if not all, of the anammox bacteria in the global ODZs. Beyond the core anammox metabolism, both organisms contain cyanase and the more dominant one encodes a urease, indicating ODZ anammox bacteria can utilize cyanate and urea in addition to ammonium. The first ODZ *Scalindua* likely derived from the benthos ∼200 million years ago. Compared to benthic strains of the same clade, ODZ *Scalindua* uniquely encode genes for urea utilization but lost genes related to growth arrest, flagellum synthesis, and chemotaxis, presumably for adaptation to the anoxic water column.

## Introduction

Anaerobic ammonium oxidation (anammox) is a core process in the global marine nitrogen cycle, estimated to be responsible for a high proportion of fixed nitrogen loss in the pelagic ocean^1, 2, 3, 4^ and in marine sediments^5, 6^. This process is mediated by microorganisms (i.e., anammox bacteria) in the phylum Planctomycetota. Due to inhibition by oxygen^7^ in the pelagic ocean, anammox bacteria are restricted to oxygen deficient zones (ODZs), confined regions with stably limited oxygen that cannot support functional aerobic respiration^8^. The three major ODZs are the Arabian Sea and the Eastern Tropical North and South Pacific (ETNP and ETSP, respectively). First discovered in the Black Sea^9^ and Golfo Dulce^10^, marine anammox bacteria and their biogeochemical significance have been extensively studied [e.g.,^2, 3, 11, 12, 13^]. Yet basic questions regarding the identity, origin, age, and adaptive mechanisms of anammox bacteria to the ODZ environment remain unaddressed.

Anammox bacteria obligately require ammonium, which is scarce across the global ocean. In addition, they can face strong ammonium uptake competition from cyanobacteria, which possess higher ammonium affinity than anammox bacteria and are also present in ODZs at certain depth horizons^14^. To possibly overcome this environmental disadvantage, some anammox bacteria are capable of utilizing alternative substrates such as cyanate and urea^13, 15^, two organic nitrogen compounds commonly present in the ocean^16, 17^. It is not known how widespread these metabolic potentials are among the dominant ODZ anammox bacteria.

Anammox bacteria are members of an old functional guild, with their last common ancestor dated to appear around the Great Oxidation Event (2.32–2.5 billion years ago^18^)^19^. However, the early-evolved, deep-branching lineages of anammox bacteria (i.e., *Candidatus* Bathyanammoxibiaceae) are mainly found in marine sediments and groundwater but are absent in ODZs^20^. Geochemical proxies suggest that the earliest emergence time of the modern-type ODZ is about 1.4 Ga in the Mesoproterozoic^21^. Whether anammox bacteria colonized the earliest ODZs and influenced ocean biogeochemistry ever since remains unclear due to the lack of diagnostic anammox signatures in the geological past.

The origin of anammox bacteria in the ODZs is another open question. Anammox bacteria in ODZ waters are mainly free-living ^22^ and originate either from other ODZs via ocean circulation or from benthic sediments. Considering a parcel of water spends hundreds of years to connect the geographically separated ODZs, it is unlikely that anammox bacteria can be maintained for such a long time under unfavorable oxic conditions. Instead, it is more conceivable that anammox bacteria in ODZs derive from benthic sediments with which they directly interact. Some of the dominant benthic anammox bacteria are also highly similar to ODZ *Scalinduaceae*^5^. It is conceivable that benthic sediments may have served as the seed bank of anammox bacteria and have driven the evolutionary radiation of anammox bacteria into the ODZs. However, this hypothesis has not been tested.

To address these questions, we leverage metagenome sequencing data from the global ODZs^23^ to reconstruct high-quality metagenome-assembled genomes (MAGs) of anammox bacteria. Our data suggest that the two ODZ anammox bacteria we recover are cosmopolitan in the global ODZs and represent the majority, if not all, of the anammox bacteria in this habitat. Their close relatives are also broadly found in marine sediments of different regions. This unique genome dataset allows us to address questions related to the identities, metabolic functions, origin, emergence time, and adaptive mechanisms of anammox bacteria in the global ODZ environment.

## Results and Discussion

### Two novel *Scalindua* anammox genomes from the Arabian Sea ODZ

We used metagenome sequencing data from the global ODZs to recover anammox bacterial genomes. We obtained two MAGs that contain distinct scaffolds (Fig. S1). The two MAGs share an average amino acid identity (AAI) of only 68.2%, suggesting that they represent two different species^24^. ODZ_A is estimated to be 95% complete with 2.3% redundancy (composed of 40 scaffolds), while ODZ_B is 92% complete with 2.3% redundancy (composed of 87 scaffolds) (Table 1). ODZ_A contains a near full-length (1,569 bp) 16S rRNA gene sequence, whereas ODZ_B does not contain a 16S rRNA gene. These MAGs of high completion and low redundancy levels enable us to perform genome-based analyses with high confidence.

To pinpoint the identities and phylogenetic affiliations of the two ODZ anammox bacterial genomes, we analyzed two sets of phylogenetic markers: (i) 120 bacterial single-copy genes (Fig. 1A) and (ii) the 16S rRNA gene (Fig. 1B). Both analyses suggest that both ODZ MAGs are novel members of the canonical *Scalindua* genus in the *Ca.* Scalinduaceae family. The canonical *Scalindua* genus is notably different from the two deep-branching genera (i.e., g Scalindua_A and g SCAELE01 in GTDB) that are mainly comprised by marine sediment representatives (Fig. 1). The deep-branching sediment members indeed have likely been erroneously named, and we propose two new genus names (i.e., *Ca.* Mariniscalindua and *Ca.* Sediminiscalindua) to distinguish them from the canonical *Scalindua* genus (Supplementary Text 1).

**Fig. 1.**
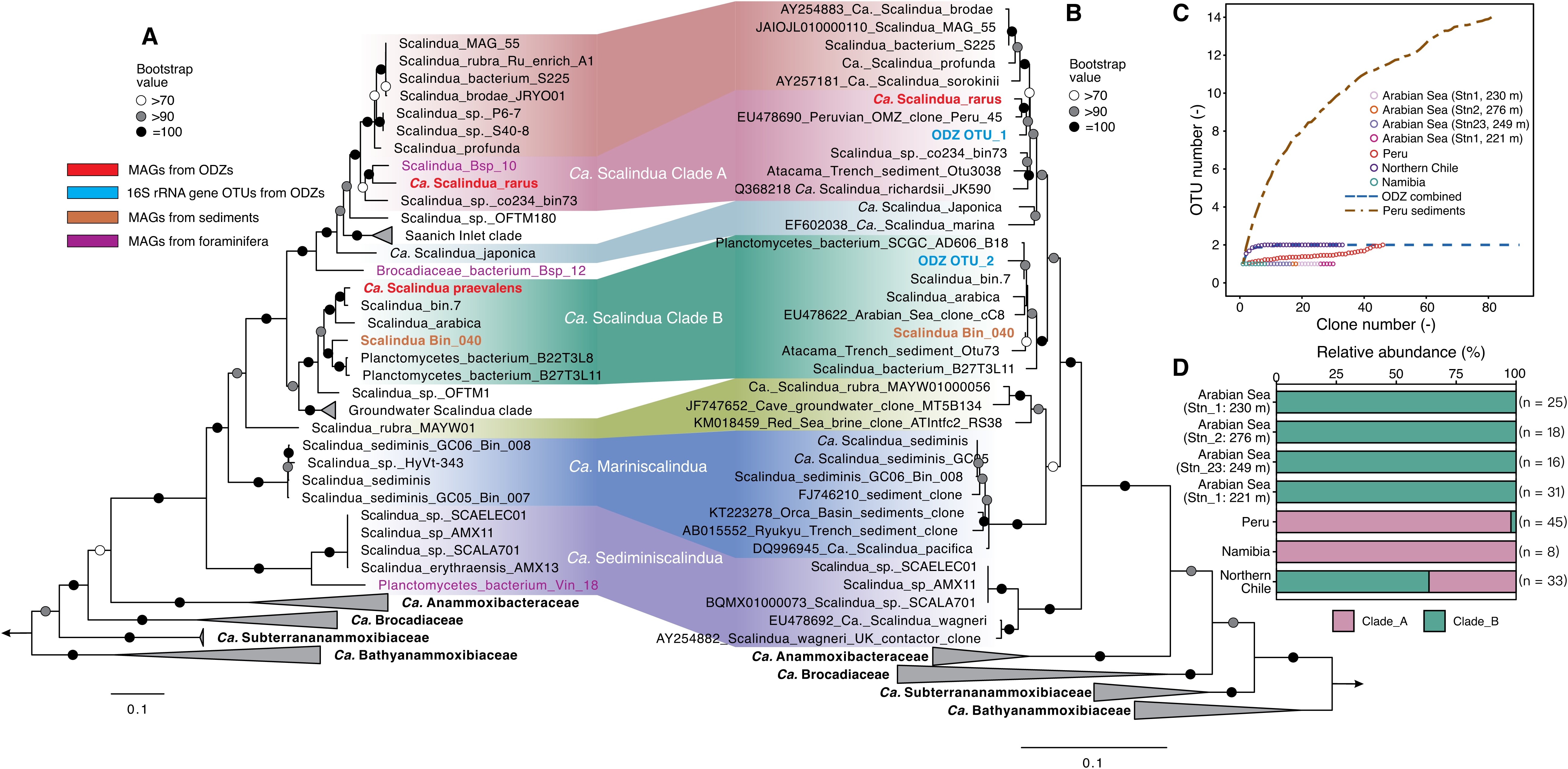
Identities and diversity of Anammox bacteria in ODZs. (**A**) Maximum-likelihood phylogenetic tree of anammox bacteria based on 120 bacterial single-copy genes. (**B**) Maximum-likelihood phylogenetic tree of anammox bacteria based on the 16S rRNA gene. In both (A) and (B), only sequences in the *Ca.* Scalinduaceae family are shown, while the other four anammox bacteria families are collapsed for readability. Bootstrap values of >70 (*n* = 1000) are shown with symbols listed in the legend. The scale bars show estimated sequence substitutions per residue. The metagenome-assembled genomes recovered from the Arabian Sea are highlighted in red, while that from AMOR sediments is shown in brown. (**C**) Low diversity of *Scalindua* community in global ODZs (including Arabian Sea, Peru margin, Namibia, and Northern Chile), as demonstrated by the rarefaction plot of only ≤2 observed OTUs versus sequenced clones. Also shown are Peru margin sediments and the combination of all ODZs. A total of 176 sequences of ODZ anammox 16S rRNA gene were analyzed, although the rarefaction curve of the first 90 is shown. (**D**) Community structure of anammox bacteria in the oxygen deficient zones in the Arabian Sea, Peru margin, Namibia, and northern Chile. Clade A (pink) and Clade B (green) correspond to ODZ_A and ODZ_B, respectively, as shown in (**A** and **B**). Panels (**C**) and (**D**) are based on a re-analysis of the clone libraries of 16S rRNA gene sequences presented in ^31, 32, 33, 52^.

Among the previously identified species, ODZ_A shows the closest relationship with *Scalindua* sp. co234_bin73 (Fig. 1A), a MAG recovered from anoxic waters of Saanich Inlet^25^ and currently the only member of the species *Scalindua* sp018648405 in the GTDB database. The calculated AAI between them is 86.9% (Fig. S3), suggesting that *Scalindua* ODZ_A represents a distinct species. Similarly, ODZ_B forms a cluster with several other MAGs from marine environments (Fig. 1A), including SRR1509794_bin.7 recovered from the ODZ core of ETNP (125 m depth at 18.9°N 104.5°W) based on existing metagenome sequencing data^26^, and *Ca.* Scalindua arabica^27^ recovered from anoxic waters of the Red Sea deep halocline. ODZ_B exhibits a 99.1% AAI with SRR1509794_bin.7 and 91.2% with *Ca.* Scalindua arabica, suggesting that ODZ_B and SRR1509794_bin.7 represent two strains of the same *Scalindua* species.

### Two cosmopolitan *Scalindua* species represent anammox bacterial diversity in ODZs

Although sourced from the Arabian Sea ODZ, we determined whether the two *Scalindua* MAGs exist in other major ODZs by recruiting reads from the existing metagenome sequencing data^23^. Our results suggest that the two *Scalindua* bacteria are present in all ODZs (Fig. 2), making them cosmopolitan across ODZs. The highest relative abundance we find is 6% of the total community in the ETNP ODZ (Fig. 2D). While seemingly up to 6% may indicate anammox bacteria are not important, they routinely represent a small fraction of the total communities^28, 29^, even though they drive the majority of fixed nitrogen loss^2^. The two *Scalindua* bacteria are confined within the ODZ cores and not detectable in oxygenated waters above or below the ODZs (Fig. 2 and Fig. S2), consistent with the premise that oxygen inhibits the anammox metabolism and therefore the growth of these specialized organisms^7, 30^. The occurrence of anammox bacteria overlaps with the accumulation of nitrite (Fig. 2), suggesting this resource remains readily available. ODZ_B is about two orders of magnitude more abundant than ODZ_A at all ODZ depths (Fig. S2), except for a single sample from the upper ODZ layer (130 m) in the Arabian Sea (Fig. 2B), suggesting that ODZ_B is the more prevalent anammox bacterium. For clarity, we provisionally name ODZ_B *Candidatus* Scalindua praevalens (prevalent in English), and ODZ_A *Candidatus* Scalindua rarus (rare in English), to highlight their contrasting abundances in the ODZs.

**Fig. 2.**
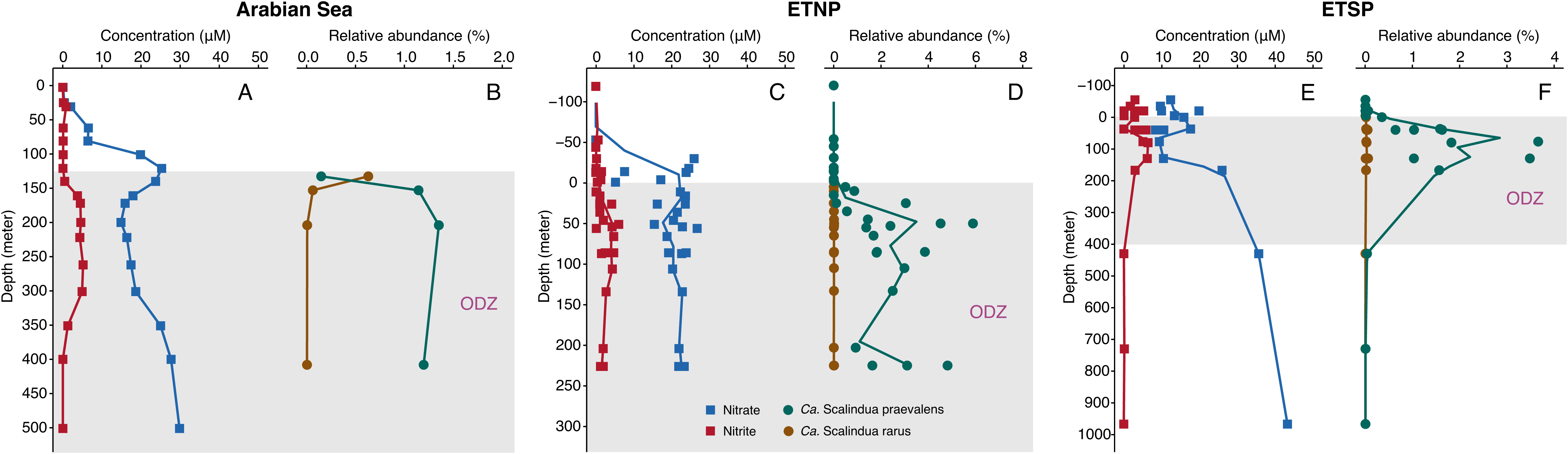
Distribution of the two *Scalindua* MAGs in three major oxygen deficient zones (ODZs). Profiles and nitrate and nitrite, and relative abundances of the two *Scalindua* bacteria in the Arabian Sea (**A, B**), Eastern Tropical North Pacific (ETNP) (**C, D**), and Eastern Tropical South Pacific (ETSP) (**E, F**). The relative abundances of the two ODZ anammox bacteria are determined by mapping multiple sets of metagenome sequencing reads onto the two *Scalindua* bacterial genomes. To integrate samples of different stations into a coherent profile for ETNP and ETSP, actual depths are converted to depths relative to the onset of the ODZ core. The circles represent individual samples, while the lines denote moving means. The waters within the ODZs are marked by a grey box at each location.

Anammox bacterial communities of extremely low diversity are characteristic of ODZs. In either individual metagenome assemblies or co-assemblies of the ODZs in the Arabian Sea and ETNP, there is maximally only one Brocadiales 16S rRNA gene sequence, which matches either *Ca.* Scalindua rarus or *Ca.* Scalindua praevalens (Table S1). Re- analyzing previous 16S rRNA gene clone library sequencing data from the Arabian Sea, Peru margin, northern Chile, and Namibia^31, 32, 33^, we find our new *Scalindua* MAGs encompass the entire diversity of the anammox community across all these sites. The total 176 anammox 16S sequences recovered from the four ODZ locations clustered into only two OTUs (operational taxonomic units, 97% nucleotide identity cutoff) (Fig. 1C). Anammox bacteria in all ODZ samples are composed exclusively of either OTU_1 or OTU_2, except that both OTUs are detected in the Peru and northern Chile ODZs (Fig. 1C and 1D). The low diversity is not likely due to primer biases (Supplementary Text 2) and is consistent with previous studies [e.g.,^28, 31^], and stands in stark contrast to the high diversity of anammox bacteria in benthic sediments (e.g., 14 OTUs are recovered via the same method from sediments underlying the Peru ODZ; Fig. 1C).

The 16S rRNA gene sequences of the two clone library OTUs match well with the *Scalindua* MAGs recovered from ODZs. Although the *Ca.* S. praevalens MAG does not contain a 16S rRNA gene, it shows 97.6% AAI with SRR1509794_bin.7, suggesting that these two MAGs represent an identical species and are interchangeable. The match between OTU_1 and *Ca.* S. rarus is evidenced by the 16S rRNA gene similarity of 98.8% over 1,526 base pairs. The same is true between OTU_2 and SRR1509794_bin.7 (i.e., *Ca.* S. praevalens), as they show 99.1% 16S rRNA gene similarity. The matches between the OTUs and ODZ *Scalindua* MAGs are supported by including the two OTUs into the 16S rRNA gene phylogenetic tree of anammox bacteria (Fig. 1B). Thus, the two *Scalindua* MAGs recovered in this study represent the most dominant, if not all, anammox bacteria diversity in the global ODZs.

### Metabolic potential of the two dominant *Scalindua* in ODZs

Similar to other characterized anammox bacteria, the two ODZ *Scalindua* have all known essential genes for the core anammox metabolism. In particular, they contain the diagnostic hydrazine synthase (HZS) for combining ammonia and nitric oxide (NO) to generate hydrazine (Fig. 3). The functional gene tree of the hydrazine synthase alpha subunit (HzsA) exhibits a similar topology to the phylogeny (Fig 1A and 1B), confirming the lineage-specific HZS in the two ODZ *Scalindua* bacteria (Fig. S3). They also have several variants of hydrazine dehydrogenase (HAO) that can oxidize hydrazine to N_2_ (Fig. 3). For the generation of NO, they can use cytochrome *cd1*-containing nitrite reductase (NirS) to reduce nitrite to NO (Fig. 3). The tree of anammox bacterial NirS (Fig. S4) shows a congruent topology with trees based on other conservative phylogenetic markers (Fig. 1), indicating that NirS is an essential trait of members of the *Ca.* Scalinduaceae family. In addition, they also contain nitrite oxidoreductase (NXR) that can provide electrons for carbon fixation through the Wood-Ljungdahl pathway (Fig. 3).

**Fig. 3.**
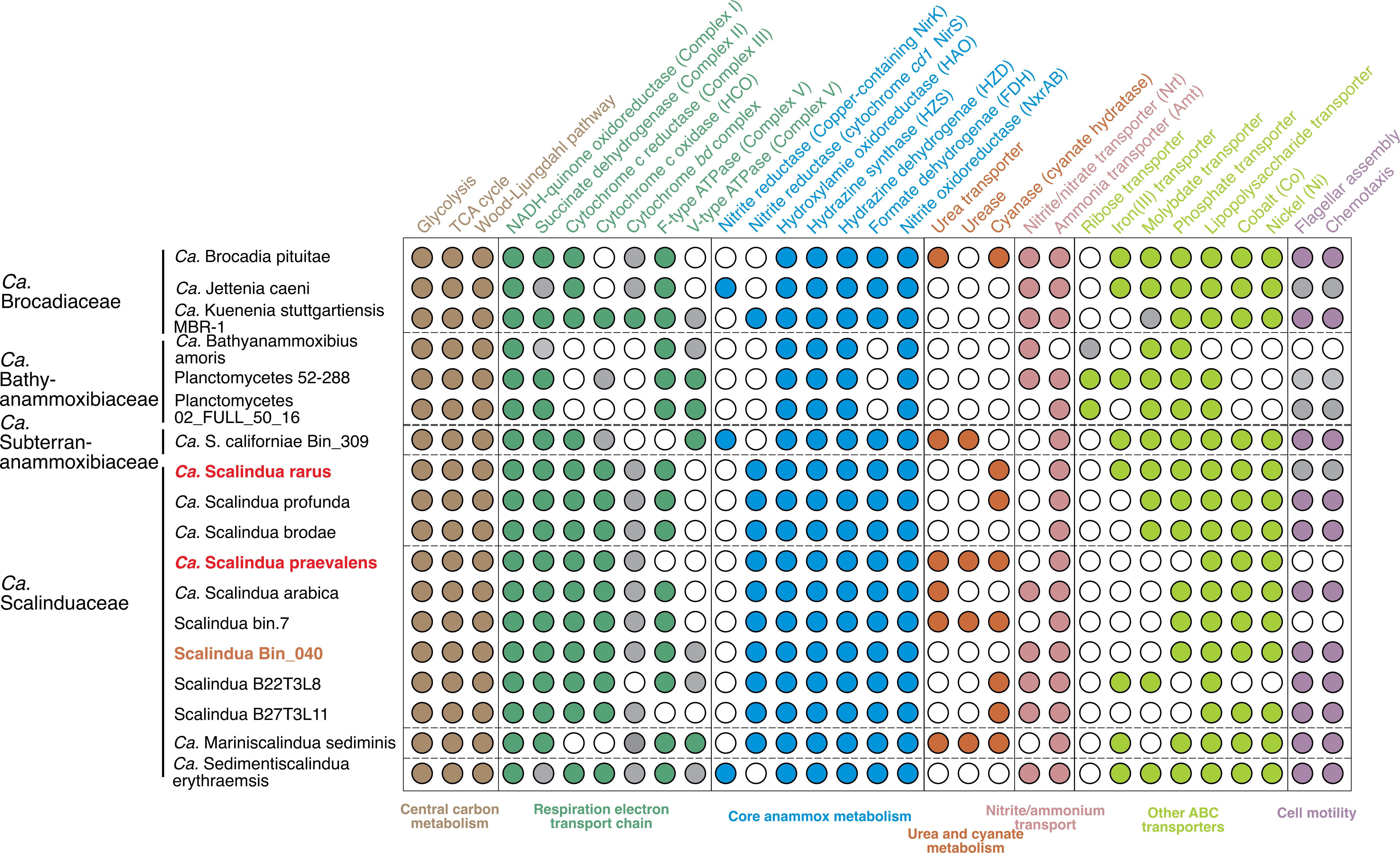
Metabolic potential encoded in anammox bacteria genomes. Filled circles indicate the presence of full pathways, open circles indicate the absence and grey ones indicate the presence of partial pathways. Genomes recovered from ODZs are shown in red, while the novel *Scalindua* genome recovered from AMOR sediments in this study is highlighted in orange. Also shown are high-quality genomes from the other three anammox bacterial families. The family-level affiliations of the anammox bacteria are noted on the left-hand side.

These high-quality *Scalindua* genomes reconstructed directly from ODZ samples provide support to their widespread capacity for utilizing diverse reduced nitrogen substrates. Both ODZ *Scalindua* bacterial genomes contain a cyanase (Fig. 3), an enzyme that can degrade cyanate to ammonium and CO_2_^34^. Given that these two *Scalindua* represent all of the known ODZ anammox bacteria diversity (Fig. 1), the presence of cyanase in both ODZ *Scalindua* bacteria indicates this metabolic trait is ubiquitous among ODZ anammox bacteria and provides direct support to the observation that cyanate stimulates anammox reaction rates^13^, presumably by providing a source of ammonium. Furthermore, this observation confirms that ODZ *Scalindua* can degrade cyanate directly. All but two cyanase-containing anammox bacteria are members of the marine *Ca.* Scalinduaceae family (either anoxic seawater or sediments) (Fig. 4). Cyanate in pelagic ODZs is produced by organic matter degradation and phytoplankton release^35^ and maintained at low concentrations due to highly active uptake relative to urea and ammonium in ODZs^16, 17^. The active expression of anammox cyanase genes^15, 36^ suggests their role in consuming available cyanate.

**Fig. 4.**
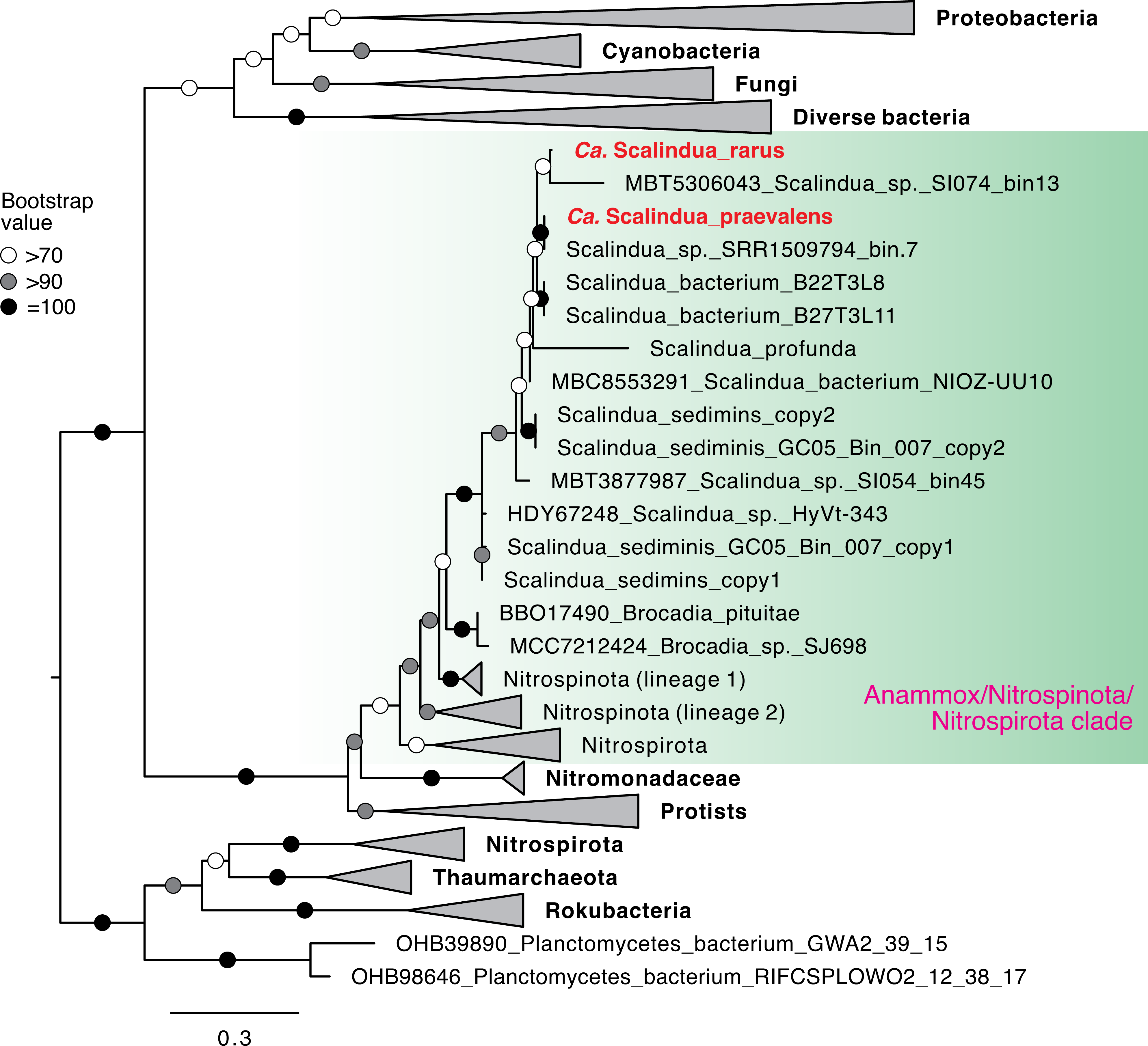
Maximum-likelihood phylogenetic tree of cyanase in anammox bacteria. For simplicity, only sequences of the clade of marine anammox/Nitrospinota/Nitrospirota clade are shown, while other clades are collapsed. The two ODZ anammox MAGs are highlighted in red. Bootstrap values of >70 (*n* = 1000) are shown with symbols listed in the legend. The scale bar shows estimated sequence substitutions per residue.

Except for *Ca.* Mariniscalindua sediminis which contains two cyanase variants^37^, all other anammox bacteria contain only one (Fig. 4). Phylogenetically, the cyanase sequences of anammox bacteria form a monophyletic clade that clusters with nitrite-oxidizing bacteria from Nitrospirota and Nitrospinota, two phyla distinct from the anammox-harboring Planctomycetota. This indicates that cyanases in anammox bacteria and nitrite-oxidizing bacteria likely result from horizontal gene transfer (HGT) events (Fig. 4). Phylogenetic similarity (inferring HGT) between anammox bacteria and nitrite-oxidizing Nitrospirota and Nitrospinota has been observed for NXR^38, 39, 40^. Cyanase is potentially another functional apparatus that has been transferred between these two important nitrogen cycling groups and may partly govern the ecological interactions between them.

Urea too is an important substrate of nitrogen-cycling organisms in the ocean^15, 41^, which mainly originates from the metabolic activities of marine organisms such as zooplankton excretion^42^ and cellular decomposition^43^. *Ca.* S. praevalens contains both a urease and urea transporter (Fig. 3). Given the abundance of this organism across ODZs (Fig. 2), it is likely that many ODZ anammox bacteria can assimilate and hydrolyze urea and produce intracellular ammonium. This observation provides strong support for the previous observation that urea can enhance anammox rates in the ETSP ODZ^13^. Urea concentrations of up to 2 µM have been detected in both oxic and anoxic seawater^17, 27^, while concentrations of ammonium in ODZs waters are only on the order of tens to hundreds of nanomolar^13, 44^, indicating that urea can be important for ODZ anammox bacteria when ammonium is limiting. In contrast to cyanate, urease is not widespread in anammox bacteria or the *Ca.* Scalinduaceae family (Fig. 3). Urease-containing anammox bacteria are only found in *Ca.* S. praevalens, several planktonic *Scalindua* genomes from Saanich Inlet^25^, *Ca.* Mariniscalindua sediminis from marine sediments, and *Ca.* Subterrananammoxibius^38^ (Fig. 3). Phylogenetic analysis of anammox urease sequences reveals that they form a clade separate from other nitrogen-cycling groups (Fig. S5), and thus urease may have a different evolutionary route than cyanase and NXR among anammox bacteria.

### Radiation of *Scalindua* into ODZs between the Carboniferous and Cretaceous

To assess for how long the two ODZ *Scalindua* bacteria have influenced ocean chemistry over geological history, we performed a molecular clock analysis to estimate the earliest divergence time of their ancestors. Although the appearance time of the last common ancestors of all anammox bacteria was previously dated to be 2.32–2.50 Ga^19^, the evolutionary history of *Ca*. Scalinduaceae members, especially the subset from ODZs, have not yet been constrained. Our analysis suggests that the origins of the clades containing the two ODZ *Scalindua* bacteria broadly fall into the Phanerozoic eon (<540 Ma, Fig. 5). In particular, the origin of the crown group of the clade containing *Ca.* Scalindua praevalens is dated to 210 Ma (95% highest posterior density (HPD) interval, 120–310 Ma), while that of *Ca.* S. rarus is dated to 190 Ma (95% HPD interval, 110–270 Ma) (Fig. 5). The origins of these two ODZ *Scalindua* bacteria are constrained in the range between the Carboniferous and Cretaceous (Fig. 5), a time when atmospheric oxygen levels were fluctuating but close to present level^45^. Considering that the earliest modern-type ODZ structure can be traced much earlier to the Mesoproterozoic [∼1.4 Ga^21^], the two *Scalindua* bacteria dominating the modern ODZs are much younger. Their appearances in the ancient oceans were at least one billion years later than the emergence of their ideal niche (i.e., minimal oxygen but sufficient supply of both oxidized and reduced nitrogen compounds) in the ocean. Our divergence time constraints raise the question of whether anammox bacteria were present and played a critical role in nitrogen cycling in the first billion years after the emergence of ODZ structure (i.e., 1.4–0.3 Ga). If yes, their niche was filled by extinct organisms. Otherwise, the ODZ nitrogen consumption during this period was likely mediated by only denitrifiers and ammonium accumulated. Nevertheless, our molecular clock analysis results suggest that the two *Scalindua* bacteria have been in the ODZs and therefore influencing global biogeochemical cycles for less than 310 million years.

**Fig. 5.**
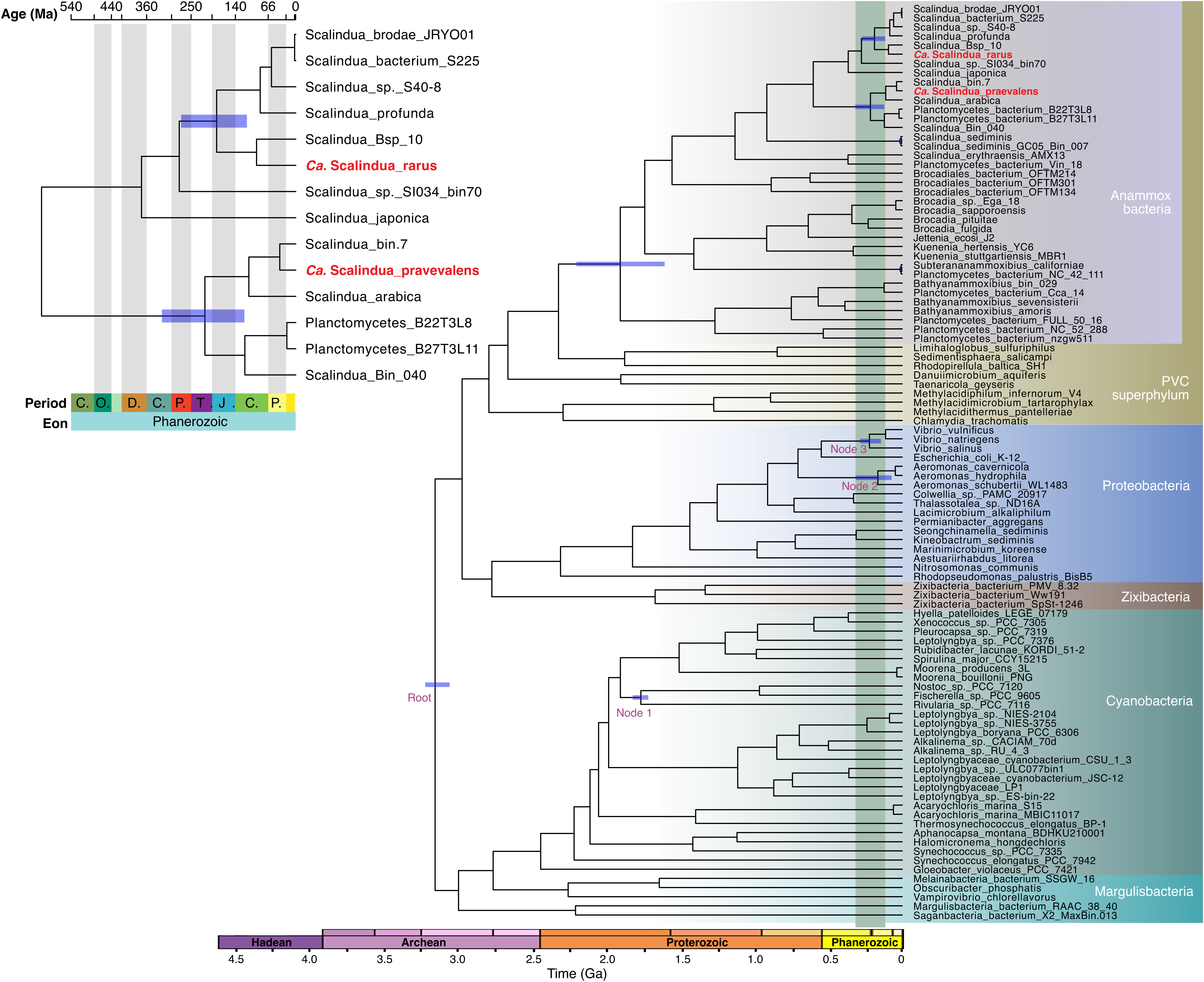
Molecular clock dating the divergence times of the two ODZ *Scalindua* bacteria. The depicted chronogram is a time-calibrated species tree that was initially generated by a concatenated alignment of 26 single-copy bacterial genes. The tree root and three nodes for calibrations are labeled in purple. Posterior distributions were generated by sampling the key Markov chain Monte Carlo analysis every 1,000 generations, with a 25% burn-in. Blue bars show uncertainty (95% confidence interval). For simplicity, only the blue bars for the nodes discussed in the main text are shown. The inset highlights the divergence times of the two ODZ *Scalindua* bacteria within the Phanerozoic eon.

### Sedimentary origin of the *Scalindua* species in modern ODZs

Despite the surprising similarity of anammox bacterial communities among the three major ODZs (Fig. 2C and 2D), we posit that direct exchanges of anammox bacteria between ODZs are unlikely. Anammox bacteria in the modern ocean are confined within ODZs^46^ that occupy only 0.1–1% of the global oceanic volume and are separated by vast oxygenated regions that effectively inhibit the anammox metabolism^47, 48^. Instead, given the abundance^37^ and relatively high diversity (containing three families) of anammox bacteria in global benthic sediments^20, 37, 38, 49^, we hypothesize that the *Scalindua* bacteria residing in modern ODZ waters derived from the benthos (Fig. 6). Material transport of metals like Fe and Mn from benthic sediments to ODZs have been previously revealed by geochemical analyses [e.g.,^50, 51^], but microbial exchanges between these two habitats are elusive.

**Fig. 6.**
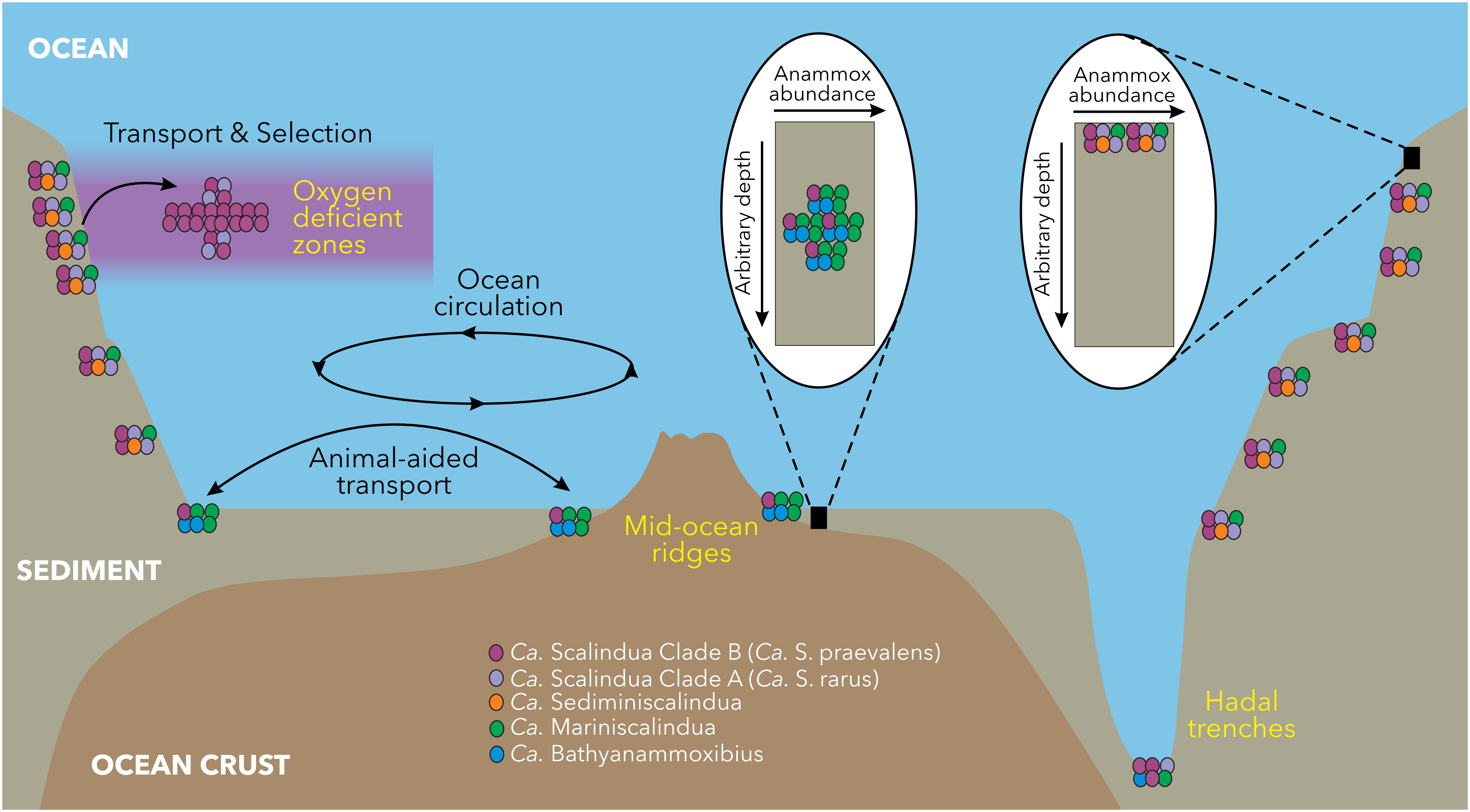
Anammox bacteria diversity across the marine environment. Anammox bacteria species are represented by different colors (not quantitative). Oxygen deficient zone anammox bacteria likely derive from benthic sediments with selection by local conditions. Anammox bacteria are widespread on the continental shelf, at mid-ocean ridges, and within hadal trenches. Because the overlying water columns in the mid-ocean ridge and hadal trenches contain no anoxic waters, the anammox bacteria found there must be transported by seawater circulation, which may be facilitated by the protection from animals such as foraminifera. In sediments, diverse anammox bacteria dominate their ideal subsurface niche, the subsurface nitrate-ammonium transition zone, the depth of which depends on organic matter supply and oxygen penetration.

We tested this hypothesis at the Peru margin, where the ODZ is in direct contact with sediments^50^. We re-analyzed the 16S rRNA gene clone library from seven sediment stations of the Peru margin^52^. The total 82 sequences clustered into 14 OTUs (97% nucleotide identity cutoff), indicating that the benthic sediments exhibit a much higher anammox bacterial diversity than the ODZs off Peru (Fig. 1C). Although the majority of the sediment sequences are affiliated with the genus *Scalindua,* especially Clade A, they also contain some sequences from the genera *Ca.* Mariniscalindua and *Ca.* Sediminiscalindua (Fig. S6). Importantly, however, the two most dominant sediment OTUs share >98.6% 16S rRNA gene identities with the two ODZ *Scalindua* MAGs (Table S2), suggesting that the two major ODZ *Scalindua* bacteria are also dominantly present in the local benthic sediments of the Peru margin. The same *Scalindua* species present in benthic sediments and the overlying ODZ waters off Peru Margin reflects potential microbial exchange between the two major habitats of marine anammox bacteria, but less diversity in the ODZs may arise from natural selection winnowing the community.

### Sedimentary relatives of *Ca.* Scalindua praevalens are widespread in marine sediments

To check if the Peru margin contains the only sedimentary relatives of *Ca.* Scalindua praevalens, we generated metagenome sequencing data from multiple marine sediment locations. We obtained a high-quality MAG (Bin_040) (91.4% complete with 1.6% redundancy, and also a 16S rRNA gene; Table 1) from Arctic Mid-Ocean Ridge sediments. This MAG has two close relatives (B22T3LB and B27T1L11, classified as *Scalindua* sp022570935 in GTDB) of lower completeness levels (<85%, Table 1) recovered from Mariana Trench sediments^53^ (Fig. 1A and 1B). While Bin_040 is not abundant (<0.16% of the total community) in the Arctic Mid-Ocean Ridge sediments where it originates, it represents the dominant anammox bacterium in the Atacama Trench sediments^54^ (Supplementary Text 3). Bin_040, together with the two Mariana Trench MAGs, form a lineage highly related to that of *Ca.* S. praevalens (Fig. 1A and 1B). Members of this genus are present with >0.01% relative abundances in 36 globally distributed marine sediment samples (Table S3). They are present in sediments not only beneath ODZs (ETNP and Arabian Sea), but also in areas without ODZs (e.g., West Pacific, South Atlantic Ridge, South China Sea, and Nordic Seas). Therefore, close relatives of *Ca.* S. praevalens inhibit benthic sediments of widespread locations (Fig. 6). This distribution implies continuity of the benthic assemblages so that the exchange of *Ca.* S. praevalens between the water column and benthic sediments can potentially happen globally, but the organism can only proliferate in favorable ODZ waters.

Novel transport mechanisms are perhaps required for the spread of anammox bacteria in global marine sediments. Considering that much of the global ocean is well-oxygenated, anammox bacteria have to be shielded from inhibitory oxygen during transport. Given that anammox bacteria are preferentially free-living^22^ and not enriched on particles^55^, they may use other objects as rafts to spread. Symbiosis is one possible mechanism to resist natural inhibition by gaining protection from another organism. In supporting this, we found four MAGs of anammox bacteria (Table 1) in foraminifera in the Peru margin sediments^56^. These foraminifera-associated anammox bacteria exhibit high diversity and are affiliated with three genera in two different families (Fig. 1A). In particular, Bsp_10 is closely related to *Ca.* S. rarus (Fig. 1A), suggesting that foraminifera may have served to shuttle this particular species between sediments and the ODZ (Fig. 6). Considering that the crown-group of foraminifera has been dated to appear in the Neoproterozoic [690–1150 Ma^57^], they were likely available as rafts during the radiation of the two ODZ Scalindua bacteria from benthic sediments to the water column. More work on the phylogenetic range of anammox-containing foraminifera in diverse locations is required to confirm this provocative hypothesis.

### Genes unique to the most dominant ODZ *Scalindua*

Given that different strains of the same species are present in the ODZs and marine sediments, we compared their genomes to identify any genes that may result in this apparent habitat differentiation among marine anammox bacteria. We focus on the clade harboring *Ca.* Scalindua praevalens (Clade_B) because it contains three highly related representatives of ODZs (*Ca.* S. praevalens, bin.7, and *Ca.* S. arabica) and also three of sediments (Bin_040, B22T3LB, and B27T1L11), while such data are not available for the clade harboring *Ca.* S. rarus. Despite smaller genome sizes (1.8–2.1 Mbp) than their sediment counterparts (2.6–2.8 Mbp), the ODZ genomes have some unique genes, including urease. Urease is present in two of the three ODZ genomes but none from sediments, which may enable them to exploit urea in seawater as an alternative substrate. In addition, we found that the ODZ genomes lack methylmalonyl-CoA mutase, which is present in the three sediment genomes. Methylmalonyl-CoA mutase can impair cobalamin acquisition and arrest growth^58^, which be advantageous for sedimentary microbes under extreme energy limitation^59^. Notably, the ODZ genomes also have no genes for flagellar synthesis and chemotaxis, which would otherwise enhance microbial colonization onto particles^60^. However, anammox bacteria in ODZs are free-living^22^, autotrophic, and their substrates are mainly dissolved in seawater. Avoiding attachment to particles can help them stay within their preferred niche rather than being exported deeper. Together, these genomic changes may have played important roles in enabling them to adapt to planktonic lifestyles in global modern ODZs.

## Conclusion

We documented two novel *Scalindua* bacteria that represent most, if not all, anammox bacteria in the global ODZs. The genome contents and the relative abundances of these two bacteria indicate that all ODZ ananmox bacteria can utilize cyanate and many can also use urea as reduced nitrogen substrates, underscoring the importance of these compounds as a source of ammonium and a new link in global ocean nitrogen cycling. These anammox bacterial communities maintain low diversity but high similarity between geographically separated ODZs and likely stem from benthic sediments where their close relatives have been found. The evolutionary radiations of these *Scalindua* bacteria from sediments into the ODZs are estimated to have happened less than ∼310 million years ago, about one billion years later than the earliest appearance of the chemical conditions identifiable as modern-day ODZs. The appearance of anammox bacteria in the ODZs may have resulted from their sediment-to-ODZ transport, before which, either ammonium accumulated or this niche was filled by other organisms. Compared to their sedimentary counterparts, the ODZ *Scalindua* bacteria acquired urease but also lost genes involved in growth arrest, flagellar synthesis, and chemotaxis. Our findings shed new light on the ecology and history of one of the major functional guilds that govern nitrogen availability in the global ocean.

## Methods

### Metagenome sequencing, read recruiting, re-assembly, and manual refinement

We reconstructed metagenome-assembled genomes of anammox bacteria from the deep-sequenced metagenome sequencing data of the ODZs of the Arabian Sea and ETNP^23^. All detailed procedures for sample and data collection were described previously in ref.^23^. Briefly, DNA extractions were performed with the Qiagen AllPrep DNA/RNA Mini Kit following manufacturer protocols. Sequencing was performed at the DOE Joint Genome Institute (JGI) on a NovaSeq (Illumina) resulting in paired-end reads of 151 base pairs in length. After quality control and trimming using Trimmomatic v0.39^61^, the reads were assembled using MEGAHIT v1.2.9^62^. Metagenome-assembled genomes (MAG) were generated from the assembly. The quality-controlled reads were grouped into genome bins using three automatic binners (CONCOCT v1.00^63^, Metabat2 v2.12.1^64^, and Maxbin2 v2.2.6^65^), implemented in the metaWRAP v1.3 wrapper^66^. All resulting MAGs were classified using GTDB-tk v2.3.0^67^ with the default setting.

To ensure the accuracy of putative anammox bacterial MAGs, we manually checked the genomes by reads recruitment, assembly, and re-binning. For each genome, quality-trimmed reads of the sample that showed the highest coverage of this genome were aligned onto the contigs using BBmap^68^, and the successfully aligned reads were re-assembled using SPAdes v3.12.0^69^ with the *k*-mers of 21, 33, 55, and 77. After the removal of contigs shorter than 1000 bp, the resulting contigs were visualized and manually re-binned using gbtools v2.6.0^70^. The input data for visualization and re-binning in gbtools include GC content, taxonomic assignments, and differential coverages of contigs across multiple samples. To generate these input data, the coverages of contigs in each sample were determined by mapping trimmed reads onto the contigs using BBMap v.37.61^68^. GC content of individual contigs was also calculated using BBMap v.37.61^68^. Taxonomic classifications of contigs were assigned by BLASTn^71^ according to the taxonomy of the single-copy marker genes in contigs. The quality of the resulting anammox genomes was checked using the “predict” command of CheckM2 v1.0.1^72^ with the default setting. The whole procedure (i.e., reads recruiting, re-assembly, re-binning, and genome quality check) was repeated multiple times until the quality could not be improved further.

### Relative abundance calculation for ODZ *Scalindua* bacteria

To determine the distribution of the two ODZ *Scalindua* in the global ODZs, we leveraged the existing ODZ metagenome sequencing data from the global ODZs [i.e., the Arabian Sea^23^, ETNP^23, 26, 73^, and ETSP^55, 74^]. We determined the relative abundances of these two *Scalindua* genomes in a total of 54 metagenome samples in the three major ODZs (Arabian Sea, ETNP, and ETSP), using CoverM using the flags minimap2-sr --min-read-aligned-percent 50 --min-read-percent-identity 0.95 --min-covered-fraction 0 (https://github.com/wwood/CoverM). For ETNP and ETSP, the relative abundances of *Scalindua* bacteria and nutrients (nitrite and nitrate) in samples of multiple stations collected during different cruises were plotted, by converting the depths to the relative depths to the onset of the ODZ (i.e., the depth with <3 µM of oxygen).

### Genome annotation

All anammox bacterial genomes were annotated using Prokka v1.13^75^, eggNOG^76^, and BlastKoala^77^ using the KEGG database. The functional assignments of genes of interest were confirmed using BLASTp^78^ against the NCBI RefSeq database. The metabolic pathways were reconstructed using the KEGG Mapper^79^. Average amino acid identity (AAI) between genomes was calculated using EzAAI^80^ using the default setting. We used the thresholds proposed by ref.^24^ to distinguish different species within the same genus, which share 65–95% AAI and 95–98.6% nucleotide identity of 16S rRNA gene.

### Phylogenetic analysis

To pinpoint the phylogenetic placements of the two ODZ *Scalindua* bacteria, we performed phylogenetic analyses for them together with the high-quality genomes of the Planctomycetes phylum included in the GTDB 08-RS214 Release (April 2023). The bacterial 120 single-copy genes were identified, aligned, and concatenated using GTDB-tk v2.3.0 with the “classify_wf” command. The maximum-likelihood phylogenetic tree was inferred based on this alignment using IQ-TREE v1.5.5^81^ with LG+F+R7 the best-fit model selected by ModelFinder^82^ and 1,000 ultrafast bootstrap iterations using UFBoot2^83^.

A maximum-likelihood phylogenetic tree based on 16S rRNA gene sequences was also used to support the phylogenetic placements of the ODZ *Scalindua* bacteria, by adding the sequences and their close relatives into the reference sequences compiled by ref.^38^. Sequences were aligned using MAFFT-LINSi^84^ and the maximum-likelihood phylogenetic tree was inferred using IQ-TREE v1.5.5^81^ with GTR+F+R5 as the best-fit substitution model and 1,000 ultrafast bootstraps, following the procedure described above.

For the phylogenetic analyses of the key enzymes of anammox bacteria (e.g., cyanase (CynS), urease alpha subunit (UreC), hydrazine synthase alpha subunit (HzsA), and cytochrome *cd1* nitrite reductase (NirS)), the sequences of the two ODZ *Scalindua* were used as the query in the BLASTp^78^ search in the NCBI database (>50% similarity and *E*-value of 10^-6^) to identify their close relatives. These sequences were aligned using MAFF-LINSi^84^ with reference sequences from ref.^37^. The alignments were then trimmed using trimAl^85^ in “automated” mode. Maximum likelihood phylogenetic trees were reconstructed using IQ- TREE v1.5.5^81^ with best-fit evolutionary models and and 1,000 ultrafast bootstraps.

### Comparative genomic analysis

We performed a comparative analysis of six genomes within the clade containing *Ca.* Scalindua praevalens. These genomes represent six strains of the same species. Three (*Ca.*Scalindua praevalens, *Ca.* S. arabica, and bin.7) are from ODZs and the remaining three are from marine sediments. We ran the analysis using Anvi’o v7.1^86^. All genomes were first annotated using anvi’o against the NCBI’s Clusters of Orthologous Groups (COGs)^87^. The comparative genomic analysis uses BLAST to quantify the similarity between each pair of genes, and the Markov Cluster algorithm (MCL)^88^ (with an inflation parameter of 2) to resolve clusters of homologous genes. The shared and unique genes in the two genomes were identified via the functional enrichment analysis^89^.

### Molecular clock analysis to date the divergence times of anammox bacteria

To estimate the divergence time of the two ODZ *Scalindua* bacteria from other anammox bacteria, we performed molecular clock analysis using the program MCMCTree from the PAML package (version 4.9j)^90^. This analysis required two input files: (i) a sequence alignment of the PHYLIP format and (ii) a corresponding Newick format phylogenetic tree of the to-be-dated microbial genomes and appropriate reference species. We selected 101 genomes in the analysis, including 37 selected high-quality genomes from the five known anammox bacterial families and 64 reference bacterial genomes. The phylogenetic analysis was based on a concatenated 26 single-copy genes, which are part of the set of 71 bacterial genes that show little horizontal transfer and therefore are suitable for phylogenetic analysis^91^. After identifying the individual genes in the genomes in Anvi’o v7.1^86^, the sequences of each gene were aligned with MUSCLE^92^, and then the individual alignments were concatenated. This alignment in fasta format was converted to the PHYLIP format using Clustal Omega^93^, for the subsequent MCMCTree analysis. Also based on the alignment, a maximum-likelihood phylogenetic tree was inferred using IQ-TREE v1.5.5^81^ with the best-fit LG+R6 substitution model. The resulting phylogenetic tree was rooted at desired branches using the tree manipulating tool nwkit^94^ and the total sequence number (i.e., 101) and the tree number (i.e., 1) were manually added as the top line of the Newick tree file.

With these two input files, the dating analysis was performed in MCMCTree with the approximate likelihood method^95^ and the iteration parameters burn-in: 10,000; sampling frequency: 50; number of sampling: 50,000. The analysis was run until convergence. The estimated age ranges of the following four nodes of the tree were manually set in the tree file to run the calibration: (i) the tree root marking the divergence of Cyanobacteria at 3.0 Ga; (ii) the *Nostocales* Crown age of 1.75–2.07 Ga^96^; (iii) the *Aeromonas* Crown age of 0.072 – 0.479 Ga^97^; and finally the *Vibrio* Crown age of 0.113–0.278 Ga^98^.

### Re-analysis of clone library sequencing data of ODZ waters and sediments

To check whether the two MAGs recovered from the Arabian Ocean ODZ can represent ODZs of other locations, we re-analyzed the 16S rRNA gene clone libraries of anammox bacteria in four ODZs: Arabian Sea, Off Peru, off Namibia, and Northern Chile. We downloaded the data from the NCBI database using the accession numbers provided^31, 32, 33^. A total of seven samples included in these three studies are all from ODZ cores [i.e., 221 and 230 m at Station 1 (17.00°N, 68.01°E), 276 m at Station 2 (19.80 °N, 64.60 °E), and 249 m at Station 23 (15.00 °N, 64.00 °E) of the Arabian Sea, 52 m at Station 202 (22.64°S, 14.30°E) of the Namibian ODZ, 35 m at Station 4 (12.03°S, 77.49°W) of the Peruvian ODZ, and 50 m at Station Pro2 (20.28°S, 70.28°W) of the northern Chile ODZ]. We aligned all sequences from the four ODZs using MAFFT-LINSi^84^ and ran the OTU clustering using the 97% nucleotide identity cutoff, as implemented in Mothur^99^. Representative OTU sequences were added into the backbone 16S rRNA gene sequences and aligned using MAFFT-LINSi^84^. A maximum-likelihood phylogenetic tree was inferred using IQ-TREE^81^ with GTR+F+R5 as the best-fit evolutionary model and 1,000 ultrafast bootstrap iterations.

Likewise, to understand the anammox community structure in the benthic sediments of the Peru margin, we downloaded the 16S rRNA gene clone library sequences reported by ref.^52^. The total 82 anammox bacterial sequences were aligned and OTUs were clustered at the cutoff of 97% nucleotide similarity using Mothur^99^. Representative OTU sequences were also added to the 16S rRNA gene tree by performing phylogenetic analysis using IQ-TREE^81^.

We also investigated for the presence of the sedimentary relatives of *Ca.* Scalindua praevalens by searching the distribution of GTDB species Scalindua sp022570935 in the Sandpiper database (https://sandpiper.qut.edu.au/), which employs SingleM (https://github.com/wwood/singlem) to search against all public metagenome datasets listed in the NCBI SRA database.

### Data availability

All raw sequencing data used in this study are available in the NCBI Short Reads Archive with the bioproject number PRJNA955304. The two ODZ *Scalindua* genomes are available under the bioproject number PRJNA1023948 with accession numbers JAWKDT000000000 (*Ca.* Scalindua rarus) and JAWKDU000000000 (*Ca.* Scalindua praevalens).

## Acknowledgments

We are grateful to all scientists and crew members of numerous cruises for their efforts in generating the ODZ metagenome sequencing data analyzed in this study. We thank Gregory Fournier and Yinzhao Wang for advice on molecular clock anaysis. This work was supported by the Simons Foundation grant 622065 and National Science Foundation grants OCE- 2138890 and OCE-2142998 to A.R.B. R.Z. was supported by the MIT Molina Postdoctoral Fellowship.

