## Supplementary Information for "Origin, age, and metabolisms of dominant anammox bacteria in the global oxygen deficient zones"

**Supplementary Text 1**

***Revisiting the classification and naming of anammox bacteria within the Ca. Scalinduaceae family***

Phylogenetic analyses indicate that in addition to *Ca.* Scalindua, there are two new genera within the *Ca*. Scalinduaceae family. The first is composed of strains of *Ca.* Scalindua sediminis mainly recovered from marine sediments of the Arctic Mid-Ocean Ridge^1^ and related cloned sequences from other sediment locations (Fig. 1B). We propose to rename the genus as *Ca.* Mariniscalindua (corresponding to g__Scalindua_A in GTDB) to distinguish it from the true *Scalindua* found in ODZs. The second is composed of *Ca.* Scalindua erythraensis originally enriched from coastal sediments^2^ and related strains^3^ and falls at the basal branch of the *Ca.* Scalinduacea family. We propose to name it *Ca.* Sediminiscalindua (corresponding to g__SCAELE01 in GTDB).

**Supplementary Text 2**

***The low diversity of ODZ anammox bacteria is not due to primer biases***

Although primers used in early clone libraries studies have their primer biases, these biases should not lead to the extremely low diversity (only two OTUs) of anammox bacteria in the four examined ODZs for the following reasons. (i) The primers (the combination of Planctomycetota-specific Pla46F and bacteria-general reverse primer 1037R/1492r) used in refs.^4, 5^ have a broad target — not only anammox bacteria in the Brocadiales order but also lineages of other orders within the Planctomycetota. (ii) the capacity of these primers to capture diverse anammox bacteria has been demonstrated in various environments [e.g., estuary sediments^6^, soils^7^, and deep-sea hydrothermal sediments^8^]. Thus, if additional anammox bacteria are present in the ODZs, they should have been amplified by the applied primers.

**Supplementary Text 3**

***The dominance of Bin_040 in Atacama Trench sediments***

We determined whether Bin_040 recovered from Arctic sediments is also present in the hadal sediments beneath the Atacama Trench^9^. The similarity of geochemical stratification and microbial communities has been shown previously^10^. Based on the re-analysis of the 16S rRNA gene amplicon sequencing data of nine Atacama Trench cores^10^, there are two OTUs (OTU_73 and OTU_3038) dominant in the anammox bacterial community. Across the total 152 sediment samples, OTU_73 comprises 73.8% of the anammox bacterial community while OTU_3038 accounts for 7.8%. The 16S rRNA gene of Bin_040 shows a 100% identity to OTU_73 (Fig. 1B), suggesting that Bin_040 can represent the most abundant *Scalindua* bacteria in the Atacama Trench sediments. This is consistent with the previous description that anammox bacteria in Atacama Trench sediments are most similar to some *Scalindua* bacteria inhabiting the Arabian Sea oxygen deficient zone^11^.

**Supplementary Figures**

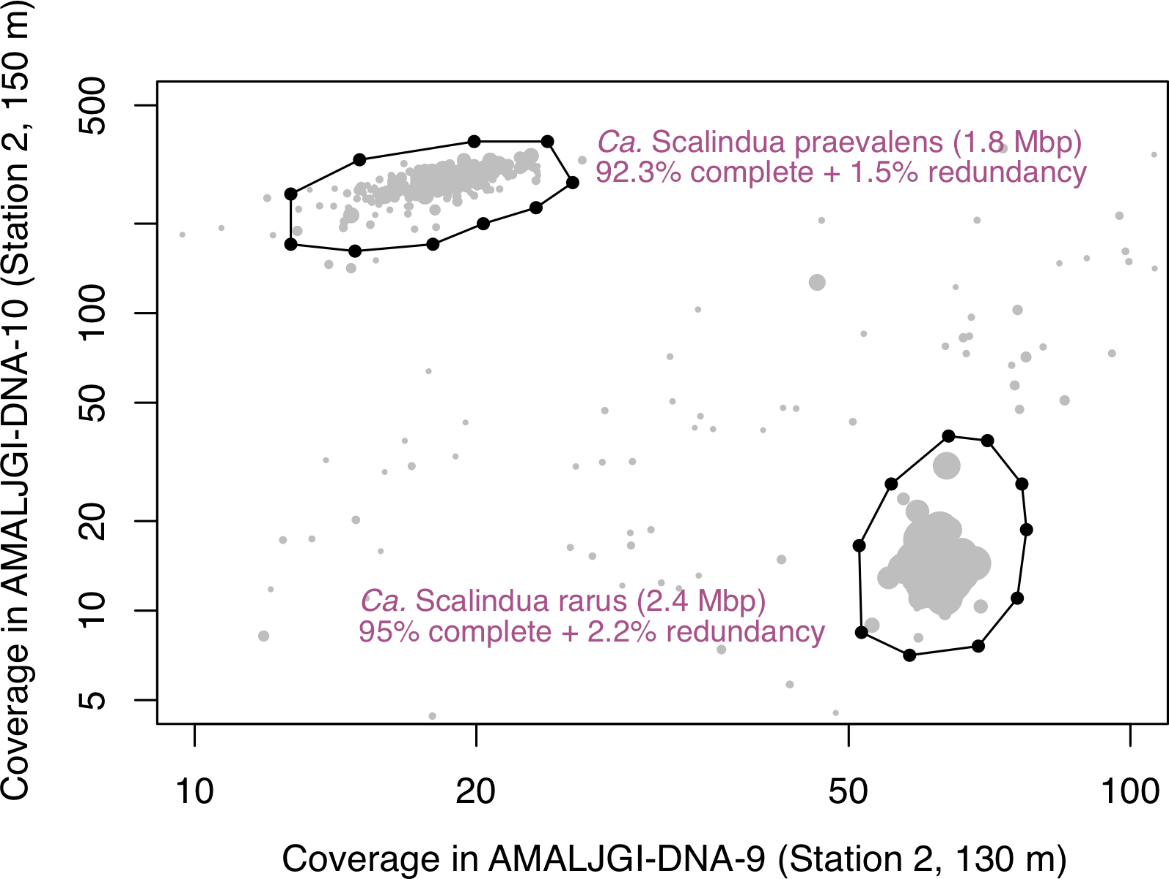

**Fig. S1. Differential coverages of the two *Scalindua* MAGs in two depths (130 m and 150 m) within the Arabian Sea oxygen deficient zone.** The scaffolds belonging to the two genomes are enclosed by two manually defined decagons.

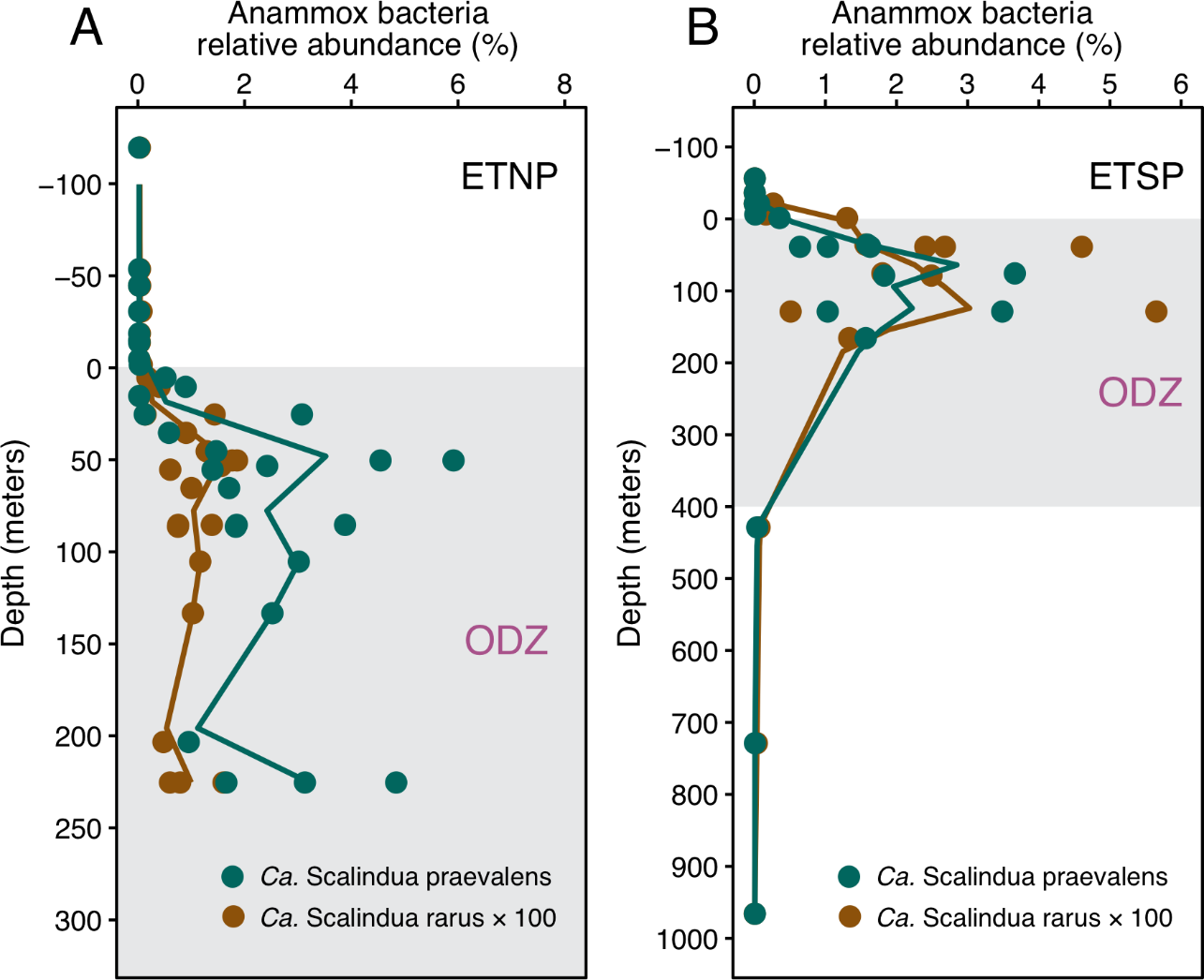

**Fig. S2. The occurrence of the two *Scalindua* bacteria in the ODZs of Eastern Tropical North Pacific (ETNP) and South Pacific (ETSP).** The relative abundances are determined by mapping multiple sets of metagenome sequencing reads onto the two *Scalindua* bacterial genomes. The depths are relative to the top of the ODZs. Circles represent individual samples, while the lines denote a running mean. The waters within the ODZs are marked by a grey box at each location. The relative abundances of *Ca.* Scalindua rarus are multiplied by a factor of 100 for visualization.

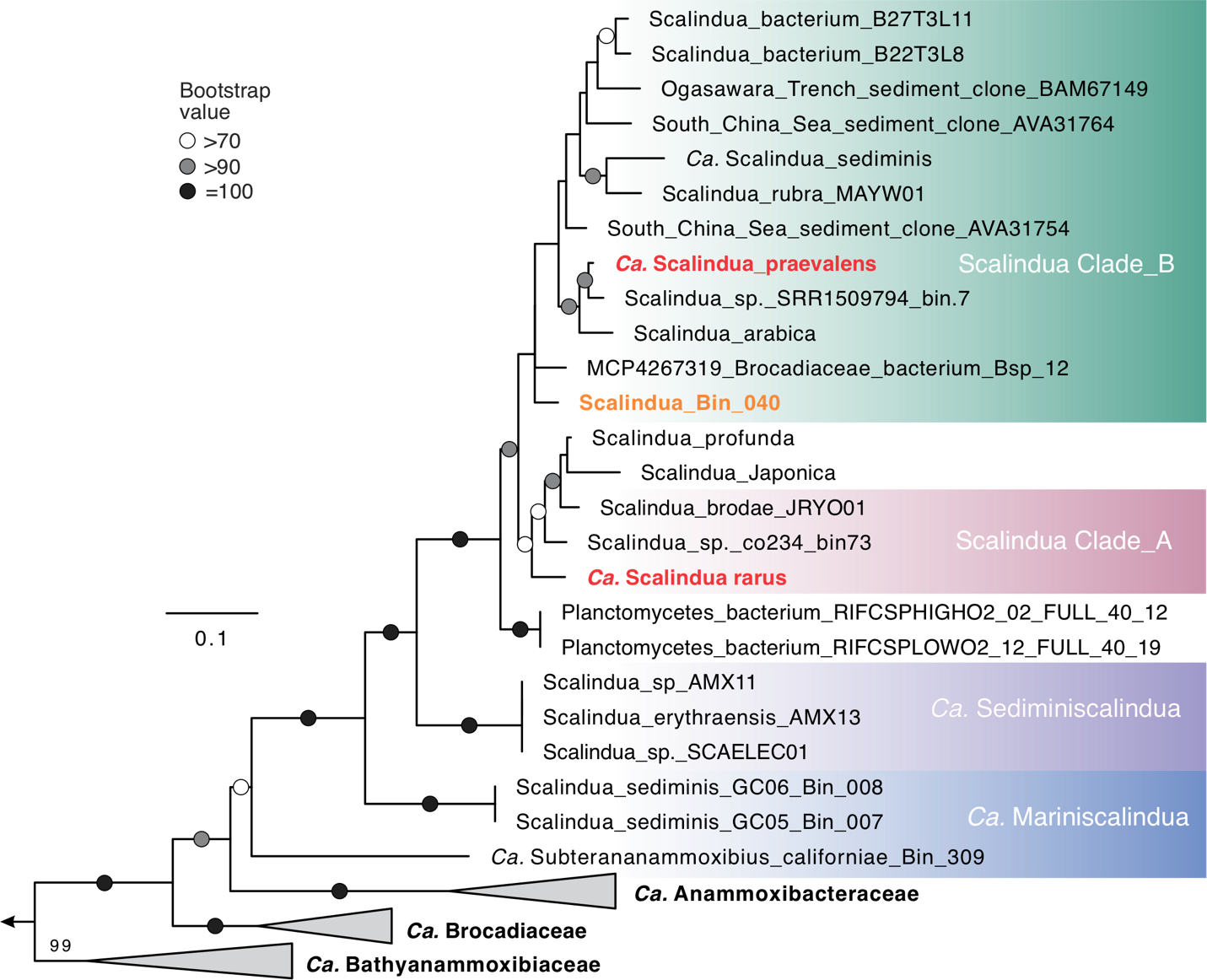

**Fig. S3. Maximum-likelihood phylogenetic tree of the hydrazine synthase alpha subunit (HzsA) of anammox bacteria.** For simplicity, only lineages within the *Ca.* Scalinduaceae family are shown, while other families are collapsed. The two ODZ MAGs are highlighted in red, while the sediment MAG is shown in orange. Bootstrap values of >70 (*n* = 1000) are shown with symbols listed in the legend. The scale bar shows estimated sequence substitutions per residue.

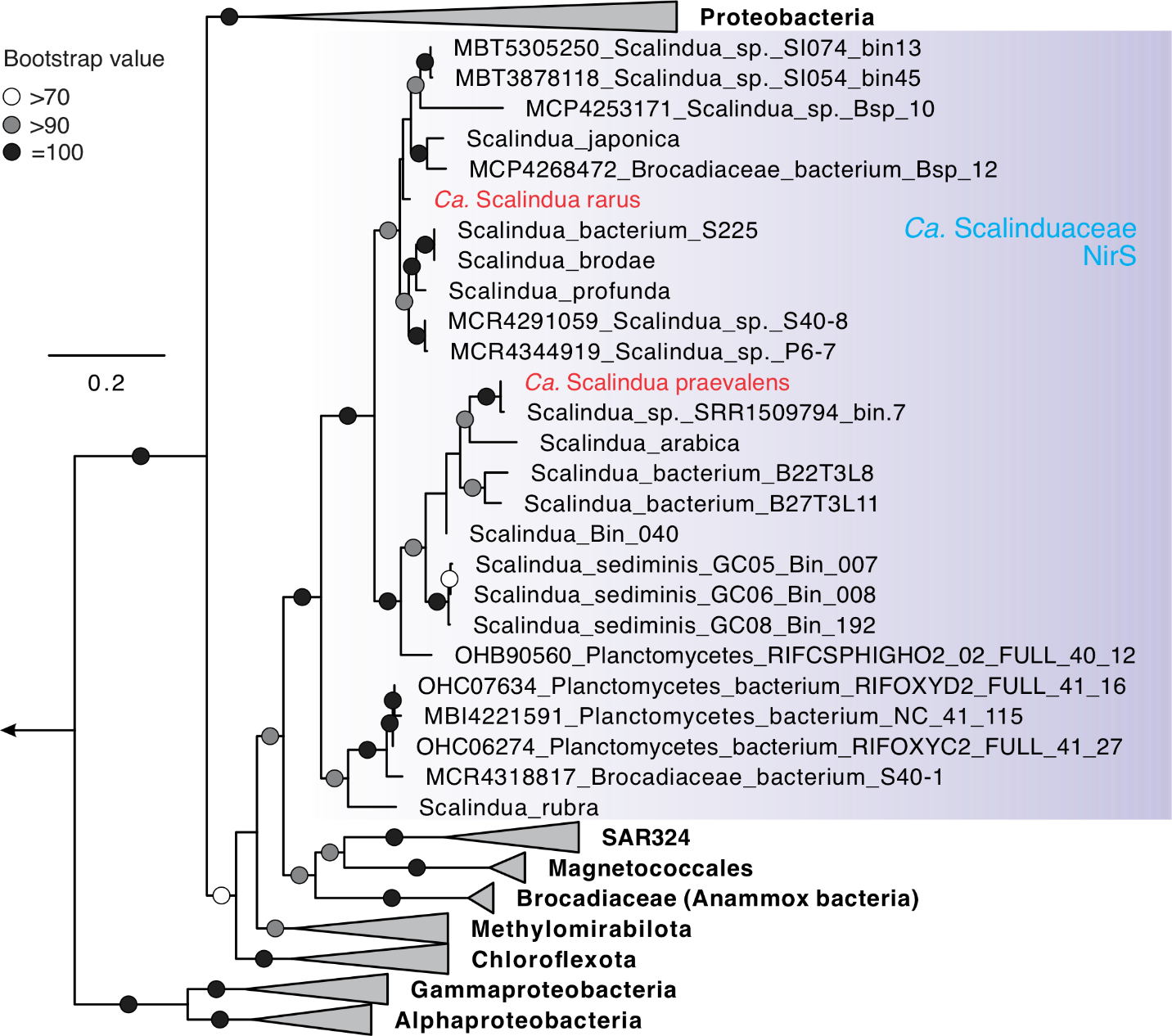

**Fig. S4. Maximum-likelihood phylogenetic tree of cytochrome *cd1* nitrite reductase (NirS) in anammox bacteria.** For simplicity, only the clade of anammox bacteria is shown, while other clades are collapsed. The two ODZ anammox MAGs are highlighted in red. Bootstrap values of >70 (*n* = 1000) are shown with symbols listed in the legend. The scale bar shows estimated sequence substitutions per residue.

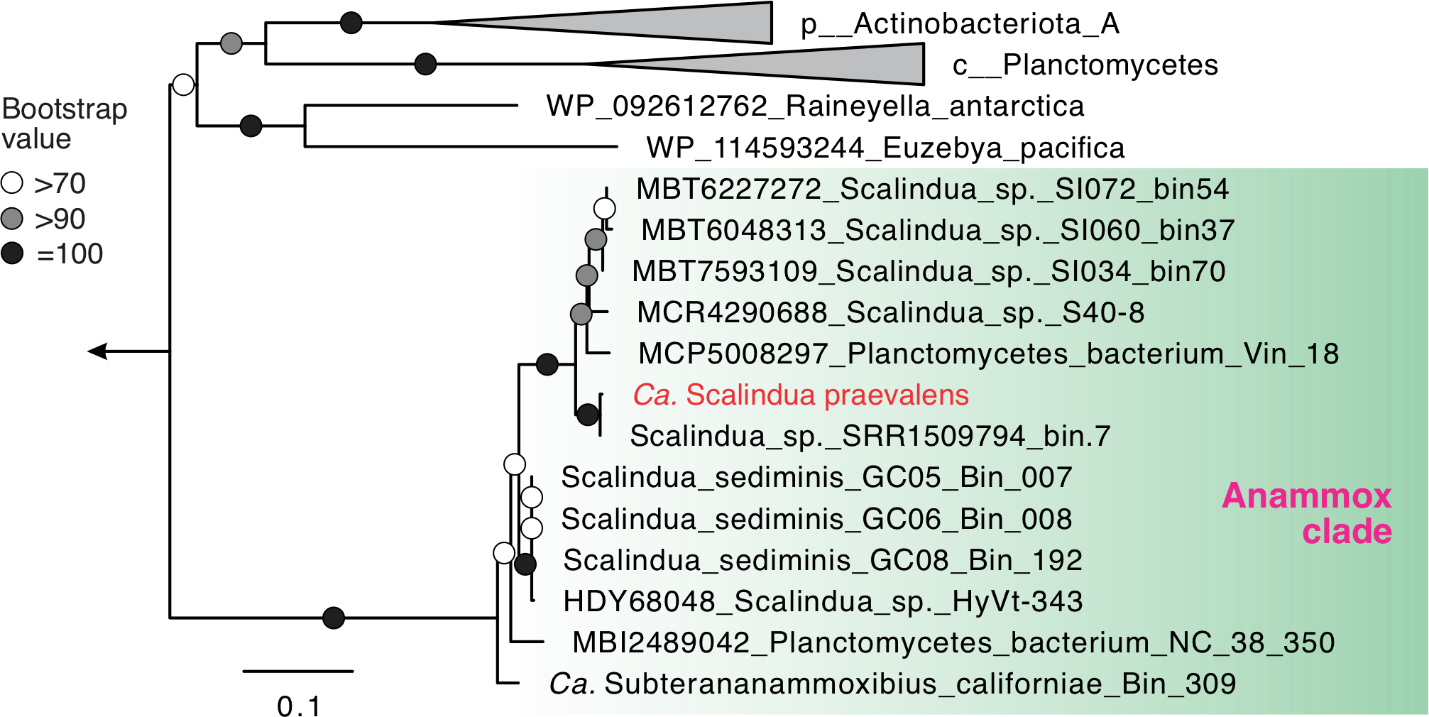

**Fig. S5. Maximum-likelihood phylogenetic tree of urease alpha subunit (UreC) in anammox bacteria.** For simplicity, only the clade of the marine anammox clade is shown, while other clades are collapsed if possible. The urease-containing ODZ anammox bacterium is highlighted in red. Bootstrap values of >70 (*n* = 1000) are shown with symbols listed in the legend. The scale bar shows estimated sequence substitutions per residue.

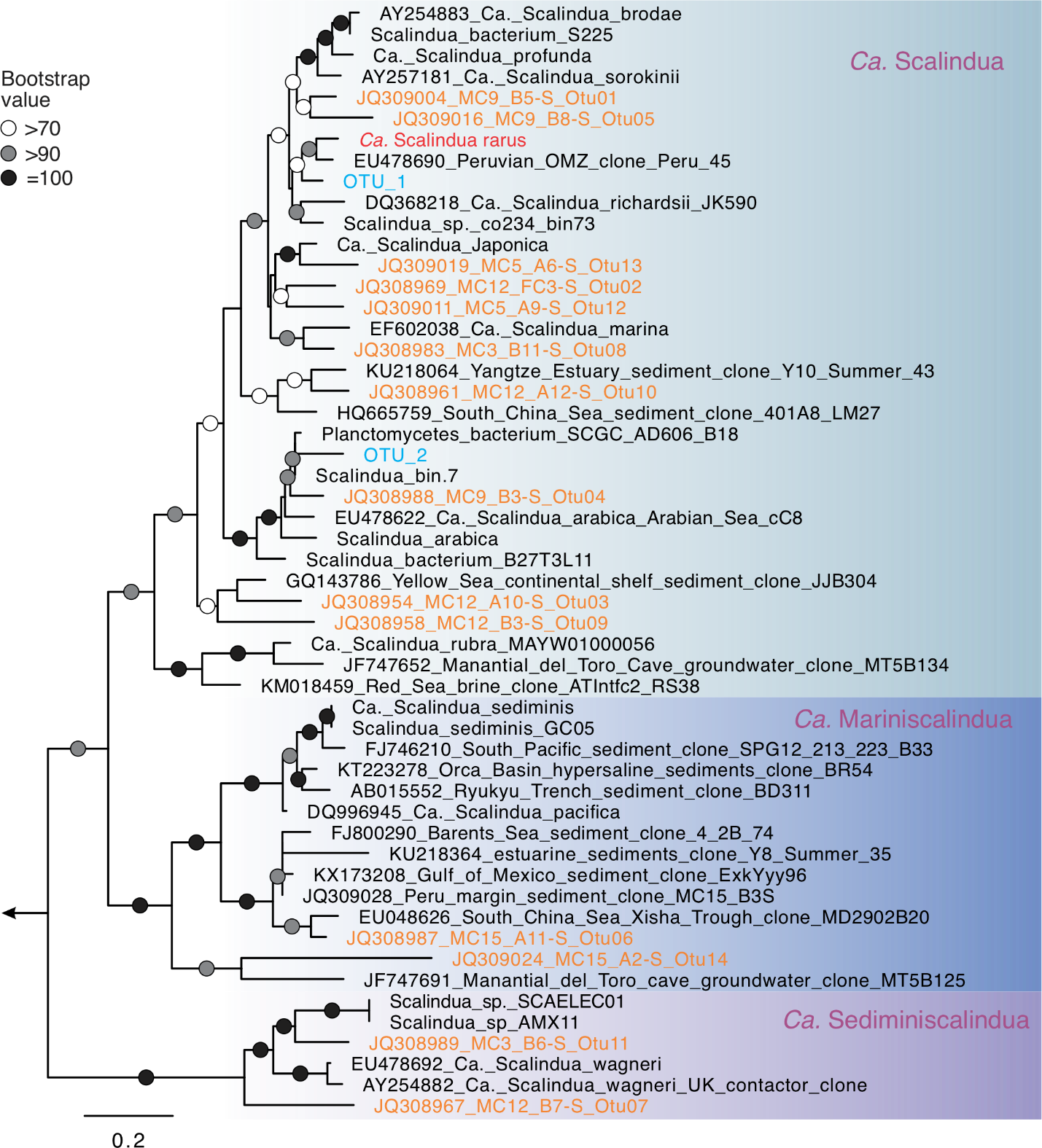

**Fig. S6. Phylogenetic affiliations of diverse *Scalindua* bacteria in sediments beneath the Peru margin oxygen deficient zone.** The sediment *Scalindua* is distributed in all three known genera in the *Ca.* Scalinduaceae family. The 14 OTU sequences from the Peru margin sediments are highlighted in orange, while the two from global ODZs are shown in blue. Bootstrap values of >70 (*n* = 1000) are shown with symbols listed in the legend. The scale bar shows estimated sequence substitutions per residue.

**Supplementary Tables**

**Table S1. Anammox bacterial 16S rRNA gene sequences in ODZ metagenome assemblies**

| Location | Metagenome SRA/JGI ID | Depth* (m) | # Brocadiales 16S rRNA  gene sequence | 16S rRNA gene phylogenetic affiliation |
| --- | --- | --- | --- | --- |
| Arabian Sea | AMALJGI-DNA-9 | 130 | 1 | *Ca.* S. rarus |
|  | AMALJGI-DNA-10 | 150 | 1 | *Ca.* S. praevalens |
|  | AMALJGI-DNA-11 | 200 | 1 | *Ca.* S. praevalens |
|  | AMALJGI-DNA-12 | 400 | 1 | *Ca.* S. praevalens |
|  | Arabian Sea co-assembly (ref. 28) | -- | 1 | *Ca.* S. praevalens |
| ETNP | AMALJGI-DNA-1 | 53 | 0 | -- |
|  | AMALJGI-DNA-2 | 120 | 1 | *Ca.* S. praevalens |
|  | AMALJGI-DNA-3 | 200 | 1 | *Ca.* S. praevalens |
|  | AMALJGI-DNA-5 | 10 | 0 |  |
|  | AMALJGI-DNA-7 | 185 | 1 | *Ca.* S. praevalens |
|  | AMALJGI-DNA-8 | 215 | 1 | *Ca.* S. praevalens |
|  | AMALJGI-DNA-13 | 60 | 0 | -- |
|  | AMALJGI-DNA-14 | 95 | 0 | -- |
|  | AMALJGI-DNA-15 | 200 | 0 | -- |
|  | AMALJGI-DNA-17 | 16 | 0 | -- |
|  | AMALJGI-DNA-18 | 45 | 0 | -- |
|  | AMALJGI-DNA-20 | 250 | 1 | *Ca.* S. praevalens |
|  | Fuchsman co-assembly (ref. 76) | -- | 1 | *Ca.* S. praevalens |
|  | Glass co-assembly (ref. 31) | -- | 1 | *Ca.* S. praevalens |

*ODZ depths are highlighted in blue, while co-assemblies are shown in yellow. --, not relevant.

**Table S2 Matches between the ODZ *Scalindua* MAGs and the dominant OTUs in Peru Margin sediments**

| Genomes | Peru sediment OTUs | 16S rRNA  gene identity | Belong to the  same species? |
| --- | --- | --- | --- |
| *Scalindua* ODZ_A | Peru_OTU_1  (JQ309004_MC9_B5-S_Otu1 in Fig. S6) | 98.6% | Yes |
| *Scalindua* ODZ_B | Peru_OTU_4  (JQ308988_MC9_B3-S_Otu4 in Fig. S6) | 98.6% | Yes |
