## Supplementary material for "Origin, age, and metabolisms of dominant anammox bacteria in the global oxygen deficient zones": Table 1

**Table 1 Summary of anammox bacterial MAGs investigated in this study**

|  | *Ca.* Scalindua rarus  (ODZ_A) | *Ca.* Scalindua praevalens (ODZ_B) | Bin_040 | MT_3L8^b^ | MT_3L11 ^b^ | Bsp_12 ^c^ | Cca_14 ^c^ | Vin_18 ^c^ | Bsp_10 ^c^ |
| --- | --- | --- | --- | --- | --- | --- | --- | --- | --- |
| Genome size (Mbp) | 2.4 | 1.8 | 2.7 | 2.7 | 2.5 | 3.0 | 2.2 | 4.3 | 3.1 |
| # Scaffolds | 40 | 87 | 255 | 616 | 270 | 410 | 66 | 711 | 432 |
| % GC | 40.1% | 40.3% | 39.6% | 39.6% | 39.2% | 38.6 | 53.1 | 41.3 | 39.3 |
| Completion ^a^ | 95.0% | 92.3% | 91.7% | 84.8% | 82.1% | 83.7% | 93.7% | 92.0% | 86.5% |
| Redundancy ^a^ | 2.2% | 1.5% | 1.7% | 2.2% | 0.8% | 5.5% | 1.1% | 5.7% | 8.5% |
| Strain heterogeneity ^a^ | 0% | 0% | 0% | 33.3% | 50.0% | 30% | 0% | 0% | 50% |
| N50 of contigs | 92,585 | 32,818 | 18,598 | 5,068 | 13,613 | 14,245 | 51,434 | 14,900 | 11,157 |
| # Coding sequences | 2,318 | 1,712 | 2,677 | 2,602 | 2,358 | 2,540 | 1,999 | 3,327 | 2,655 |
| Coding density | 86.7% | 88.7% | 86.2% | 81.6% | 80.8% | 82.7% | 87.9% | 82.9% | 83.0% |
| rRNAs | 3 | 3 | 4 | 1 | 3 | 1 | 0 | 0 | 8 |
| tRNAs | 40 | 41 | 38 | 30 | 31 | 30 | 40 | 39 | 34 |

^a^ Estimated by CheckM2;

^b^ MAGs reported by (Zhou et al., 2022); the full IDs are B22T3LB and B27T1L11.

^c^ MAGs refined based on data from (Woehle et al., 2022).

Woehle, C., Roy, A.-S., Glock, N., Michels, J., Wein, T., Weissenbach, J. et al. (2022) Denitrification in foraminifera has an ancient origin and is complemented by associated bacteria. *Proceedings of the National Academy of Sciences* **119**: e2200198119.

Zhou, Y.-L., Mara, P., Cui, G.-J., Edgcomb, V.P., and Wang, Y. (2022) Microbiomes in the Challenger Deep slope and bottom-axis sediments. *Nature Communications* **13**: 1515.
